# Kinematic signatures of fast feedback trajectory control during goal-directed finger, arm, and saccadic eye movements

**DOI:** 10.1101/2023.08.09.552575

**Authors:** Niranjan Chakrabhavi, Varsha Vasudevan, Varadhan SKM, Ashitava Ghosal, Aditya Murthy

**Affiliations:** Centre for Neuroscience, Indian Institute of Science, Bengaluru 560012, India; Department of Bioengineering, Indian Institute of Science, Bengaluru, 560012, India; Department of Applied Mechanics, Indian Institute of Technology Madras, Chennai 600036, India; Department of Mechanical Engineering, Indian Institute of Science, Bengaluru 560012, India

**Keywords:** motor control, trajectory planning, variability, and decorrelation

## Abstract

Goal-directed eye, hand, and finger movements follow invariant kinematics consisting of approximate straight-line trajectories and bell-shaped velocity profiles. A fundamental unresolved issue is whether these trajectories are planned or whether they are a consequence of trajectory-free online control. We address this question using Spearman’s rank correlation, zero-crossing rate, and z-scores, and analyze within-trial variability to investigate differences in the time evolution of trajectories during the presence or absence of a goal in finger and whole-arm reaching movements, along with analyzing rapid goal-directed saccadic eye movements. We found that the central nervous system (CNS) implements control to follow an average trajectory, where goal-directed movements show an enhanced degree of trajectory control. Further, we found behavioral signatures of rapid control that might operate on these planned trajectories as early as ∼60 ms in finger movements and ∼70 ms in whole-arm reaching movements. Such early signatures of control suggest that the system could exploit internal feedback along with fast feedback processes to implement meaningful corrections during simple voluntary unperturbed movements. The analysis also revealed that the controller gains varied along the movement and peaked distinctly at early (20 %) and late (90 %) phases of finger and arm movements, implying that trajectory control may be accomplished through implicit way-point objectives during the execution of the movement. These later corrections were missing during the late phase of eye movements, pointing to a predominant role of delayed sensory feedback mediating corrections in the context of finger and arm movements during this period.

## Introduction

Normative goal-directed movements are characterized by invariant-kinematic patterns that consist of straight-line trajectories with bell-shaped velocity profiles that scale with movement amplitude and speed (Morasso, 1981). Numerous psychophysical and neurophysiological studies highlight the notion that movement attributes are represented/programmed in the central nervous system well before execution (Keele, 1968; Rosenbaum, 1985; Dick *et al*., 1986; Morris *et al*., 1994; Moran & Schwartz, 1999; Padoa-Schioppa *et al*., 2002; Churchland *et al*., 2006; Summers & Anson, 2009). Such results have motivated the use of internal models wherein a complete specification of a trajectory can be made, in principle, before its implementation (Bizzi *et al*., 1984; Wolpert *et al*., 1995; Wolpert & Kawato, 1998; Desmurget & Grafton, 2000; Wolpert & Ghahramani, 2000). However, other influential models of motor control suggest that only certain critical features of the movement such as the target or desired displacement need to be specified in advance (Todorov & Jordan, 2002; Desmurget & Grafton, 2003; Scott, 2004); here, the neural representation specifying movement trajectories may only be generated dynamically in real-time during movement execution as a consequence of solving a cost function based on sensory feedback that is sensitive to the demands of the task. Thus, it remains unclear whether movement attributes such as trajectories are planned before movement execution or whether they are derived from online feedback control.

One feature that, in principle, can distinguish between planned and derived movement trajectories is the timing of their respective control signals that are necessary to mitigate the effects of motor noise, which is a ubiquitous property of muscle and neural activity. In the context of planned movement trajectories, a comparison between desired and estimated current states can be used as an error signal to control movements in the absence of actual sensory information (Gordon & Ghez, 1987; Kawato *et al*., 2003; Richardson *et al*., 2011). In contrast, sensory feedback control relies on relatively delayed sensory feedback to make meaningful corrections during the movement (Todorov & Jordan, 2002; Wolpert *et al*., 2011). While such rapid control has been studied in the context of fast saccadic eye movements (Kawato, 1999; Gaveau *et al*., 2003; Richardson *et al*., 2011) as well as rapid corrections to task-related errors (Cluff & Scott, 2015; Scott *et al*., 2015; LoTemplio *et al*., 2023), distinct signatures of such rapid control in the context of unperturbed normative movements have not been experimentally demonstrated to the best of our knowledge. Furthermore, in both saccadic eye movements and skeletal reaching movements, it is unclear whether these corrections are guided towards a planned trajectory (Messier & Kalaska, 1999; Richardson *et al*., 2011) and whether such trajectory corrections are constantly implemented along the movement or whether the system prefers hotspots of control during certain phases of movement (Liu & Todorov, 2007; Kuang & Gail, 2015).

We developed two novel statistical measures to quantify the extent of control/corrections to trajectories during normative goal-directed movements by investigating unique patterns among within-trial variability. Firstly, we used Spearman’s rank correlation to track the relative locations of trajectories to one another, along the duration of the movement, and associated the decrease in correlation (or the lack of) to quantify the extent of online control. This measure of control signifies online corrections independent of a planned trajectory. Secondly, we used z-scores and zero-crossing rate to quantify the propensity of trajectories to approach an average trajectory, i.e., to examine the system’s ability to follow an average planned trajectory (trajectory control). Using these statistical measures we showed presence or absence of rapid control across different movement modalities such as precise finger movements (Chakrabhavi & Skm, 2019), whole arm simple reaching movements, and across different movement contexts i.e., in the presence or absence of a goal/task, between varied skill of movements using the dominant and non-dominant arms, across different movement speeds, and during fast saccadic eye movements (Varsha *et al*., 2021).

## Materials and Methods

A total of four experiments have been conducted and analyzed for the current study, comprising a variety of finger, whole arm, and saccadic eye movements. Different task conditions have been tested for the presence or absence of control towards a planned trajectory during goal and non-goal finger movements (Chakrabhavi & Skm, 2019), goal and return movements by the arm, movements performed by dominant and non-dominant arms and at varying movement speeds (fast and slow), and saccadic eye movements in various directions (Varsha *et al*., 2021).

### Experiment 1: Cyclic flexion and extension movements in fingers

The experiment was conducted by Chakrabhavi and SKM (2019), and the data have been adopted for the current study. The participants were asked to sit on a height-adjustable chair and were made to rest their forearms and palms on a flat surface in neutral (pronation-supination) forearm posture (Fig. 1A, left). The forearm and the thumb were constrained to the experimental setup while the index, middle, ring, and little fingers were suspended in air and free to move. Electro-magnetic 3D tracking sensors (Micro Sensor 1.8^TM^, Liberty tracking system, Polhemus Inc., Colchester, VT, USA) were attached to the dorsal aspects of phalanges on all four fingers, and the data from the distal phalanges have been used in this study. The experiment was designed on the LabVIEW interface (National Instruments Corp, Austin, TX, USA) and was acquired at a 100 Hz sampling frequency. Eight healthy right-handed subjects participated in the study (5 males and 3 females), and the experiment was approved by the institute ethics committee at IIT Madras, and written consent was obtained from all the participants before the start of the experiment.

**Figure 1:**
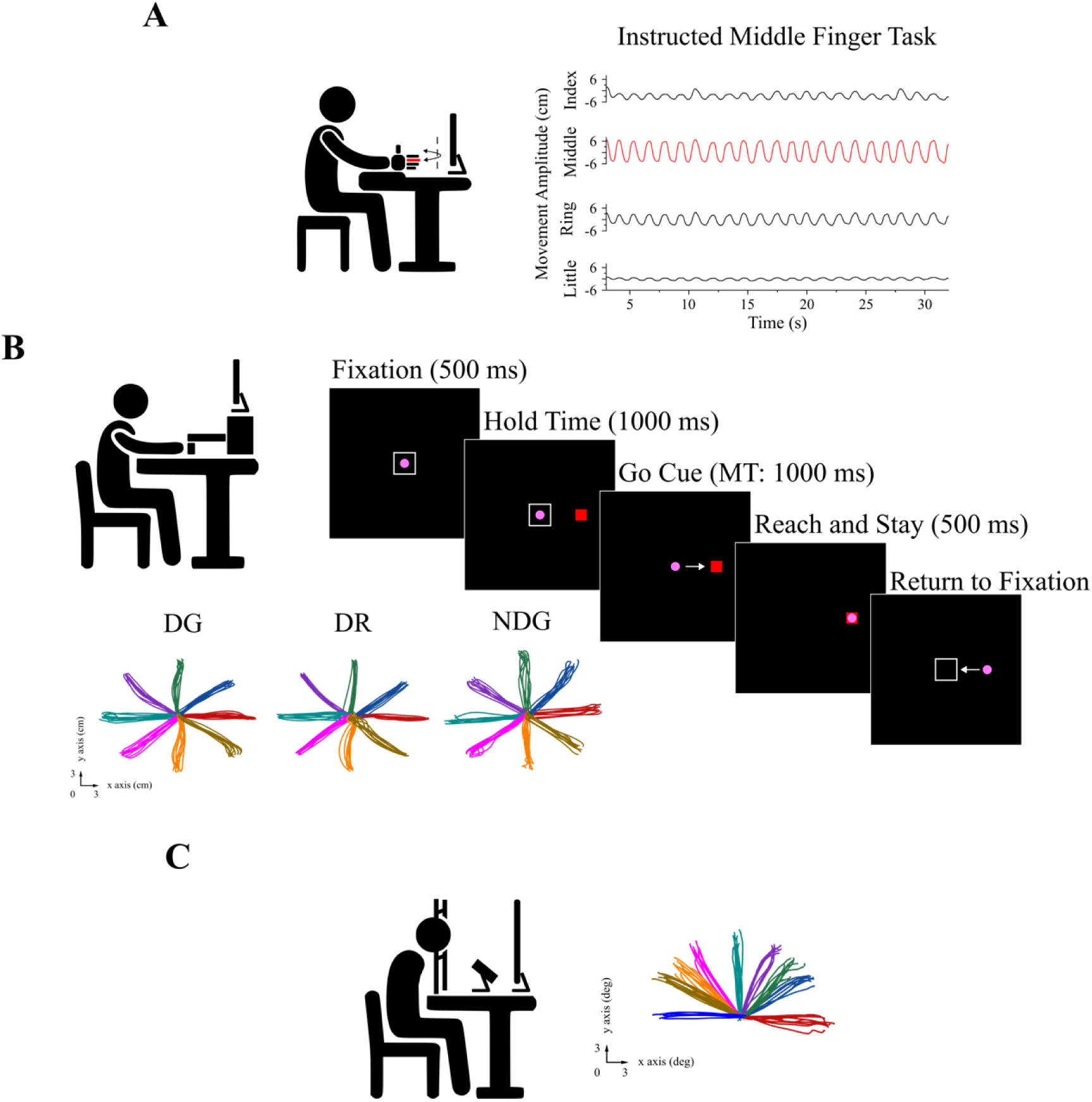
Task design of finger, arm, and saccadic eye experiments and their corresponding raw trajectories. *A) Experiment 1: Cyclic finger flexion-extension* – *Left:* The subjects rested their forearm and palms on a fixed platform in a neutral posture (pronation-supination), the thumb finger was restrained to the setup, and the four fingers (index, middle, ring, and little fingers) were suspended and free to move. On any given trial, the goal finger (middle finger, depicted with red color, in this case) was instructed to produce cyclic flexion (peaks) and extension (troughs) movements to specified landmarks shown on the stimulus screen (see methods). The phenomenon of finger enslavement led to involuntary movements in the other non-goal fingers (index, ring and little fingers). *Right:* Sample traces of goal/instructed middle finger task. The traces were from the sensors attached to the distal phalanges of the fingers. In this figure, the goal-directed middle finger movement (red color trace) engaged non-goal movements in the index, ring, and little fingers. *B) Experiment 2,3: Discrete horizontal whole arm reaching movements* – *Left:* The experimental setup consisted of a robotic arm that the subject held onto and performed simple reaching movements in the horizontal plane by looking at the stimulus presented on a screen placed in front. 3D motion trackers were mounted on the neck, shoulder, elbow, and wrist joints and on the dorsal side of the hand. The sensor on the hand was considered for the analysis in this study. *Right:* The experimental protocol started with the subject fixating a pink circular cursor in a central fixation box for 500 ms. A red-colored target appeared at an eccentricity of 12cm on the screen. The target location was block randomized to appear at any of the 8 directions (0, 45, 90, 135, 180, 225, 270, and 315 degrees from the center). Subjects had to stay at the fixation for 1000 ms during hold time, and the central fixation box disappeared, marking the go cue to begin the movement. Subjects made simple, smooth movements towards the target location within 1000ms and stayed there for an additional 500ms to complete the trial. These movements were considered goal-directed movements. They then had to get back to the central fixation box for the start of the subsequent trial. These movements were considered return movements and were not rewarded. The intertrial interval was 1500 ms. *Bottom:* Sample traces of goal (DG) and return (DR) movements by the dominant arm and the goal movements by the non-dominant arm (NDG) of one subject (10 trials each). The goal-directed movements were from the center to a target location, while the return movements were from the target to the center. *C) Experiment 4: Discrete saccadic eye movements – Left:* The subjects sat on a chair, head fixed, and performed discrete saccadic eye movements from the center to a target (randomized, 9 directions – 0^0^, 30^0^, 45^0^, 60^0^, 90^0^, 120^0^, 135^0^, 150^0^ and 180^0^) at an eccentricity of 12 degrees, while their saccadic movements were recorded using an eye tracker. The saccade had to be initiated as soon as the target appeared on the screen. *Right:* Sample trajectories of saccades along the 9 directions (10 trials each) by one sample subject.

**Figure 2:**
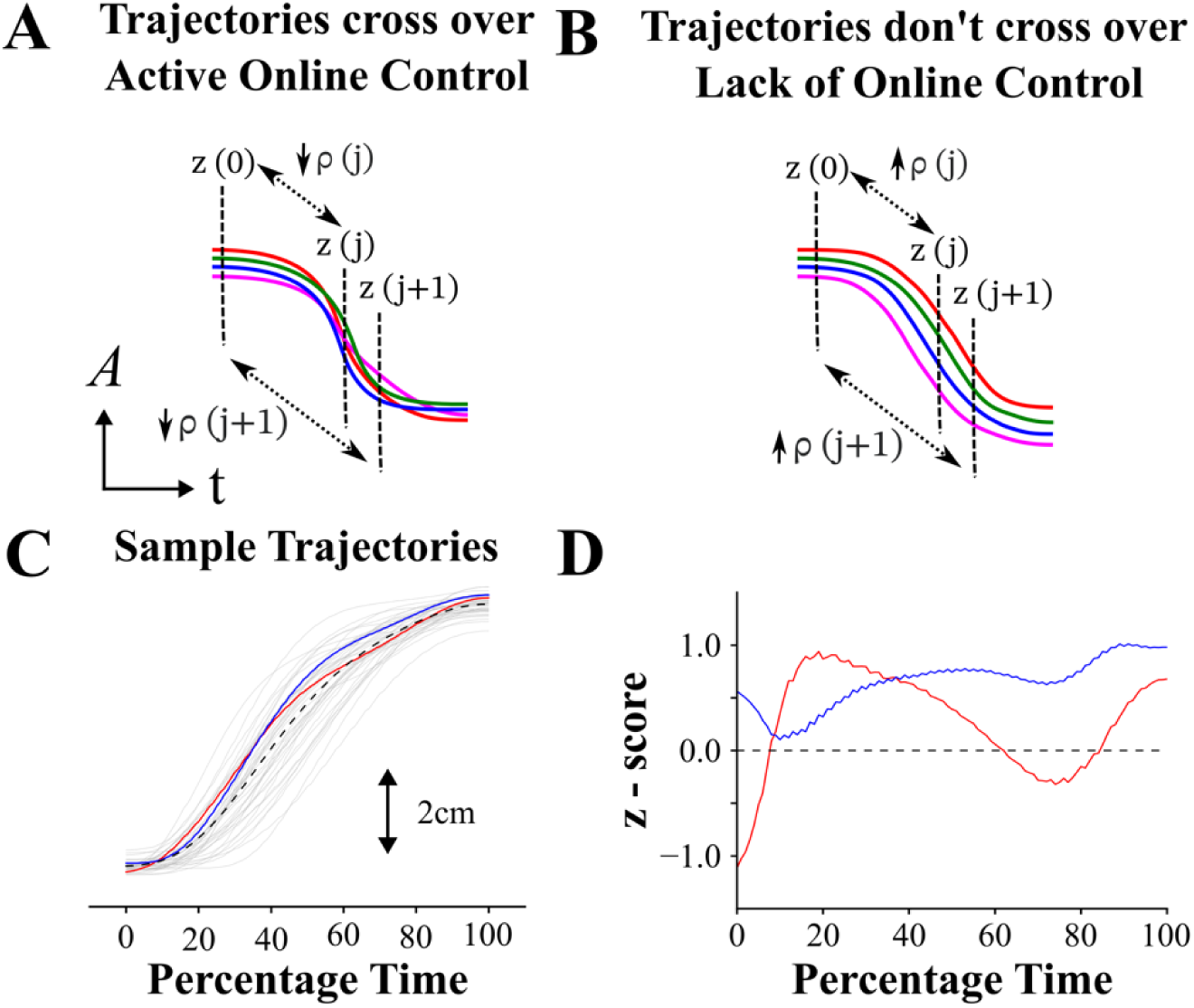
Behavioural signatures of online control and average trajectory planning. The colors red, blue, green, and pink in A and B represent individual trials of a given task. A) If the CNS implements active online control/correction based on its initial posture, the z-scores at the start of the movement z(0), are posited to change along the time course (z(j) at any time t = j), resulting in crossovers among the trajectories. These crossovers could be captured by a decrease in Spearman’s correlation coefficient (ρ). By performing the correlation analysis between movement onset and the subsequent time instances t = *0,…,j,j+1,…T* (where *T* is the end of movement), we obtain a time-varying ρ(*t*). A low value of ρ indicates greater control. B) A case where the CNS does not implement online control, i.e., the trajectories starting farther away from the average remain farther away. The correlation coefficient ρ remains high and closer to 1 along the time course. C) Sample trajectories of the X coordinate of hand movements, directed along 0^0^, of a single subject, plotted against percentage time. The dashed black line represents the average trajectory, and red and blue are two selected trials. D) Plot of *z*-scores for the red and blue trials/trajectories and the average trajectory, *z* = 0 (dashed black line), as depicted in C. To investigate if the implemented online control was directed towards an average planned trajectory, we computed the number of times the trajectories crossed the *z* = 0 line for a given set of trials (zero-crossing rate - *ZCR*, see methods). For example, the red trial has three zero-crossings during the movement and is under stronger trajectory control, i.e., a greater propensity towards the average trajectory, while the blue trial had no zero crossings and is said to be under weaker trajectory control.

During an attempt, subjects were asked to maintain a wrist posture (0^0^ neutral, 30^0^ flexion, or 30^0^ extension). They were instructed to make cyclic flexion-extension movements at the metacarpophalangeal joint with the one of the four fingers (index, middle, ring and little) between 75 ± 10 % of its maximum flexion-extension limits, and to ignore any involuntary movements in the other non-instructed fingers due to the phenomenon of finger interdependence/enslavement (Häger-Ross & Schieber, 2000; Li *et al*., 2004). The movements performed by the instructed fingers were considered goal-directed, and the involuntary movements in the non-instructed fingers were considered non-goal-directed movements. Each attempt lasted for 32 seconds, with an initial 2 seconds of rest followed by 30 seconds of cyclic flexion-extension movements (Fig. 1A, right). Real-time visual feedback of the distal phalange of the instructed finger was provided on the screen, along with its movement landmarks, and an audio metronome was played at 1.5 Hz to ensure the amplitude and timing of the instructed finger movement, respectively. There were 36 attempts (3 wrist postures x 4 instructed fingers x 3 attempts each) in total. Data from the distal phalanges of each of the four fingers at neutral wrist posture (0^0^) were used in this study. Specifically, each attempt was segmented into several half-cycles comprising movements from flexion (local peaks of middle finger trace in Fig. 1A, right) to subsequent extension (local troughs of middle finger trace in Fig. 1A, right) of instructed finger and the same time stamps were used to segment the movements of non-instructed fingers. The half cycles were then resampled to 100 equally spaced segments to analyze control with respect to a normalized percentage time scale as in Figs. 3 and 4, whereas the first 300 ms were used for the analysis on an actual time scale in Fig. 3C.

**Figure 3:**
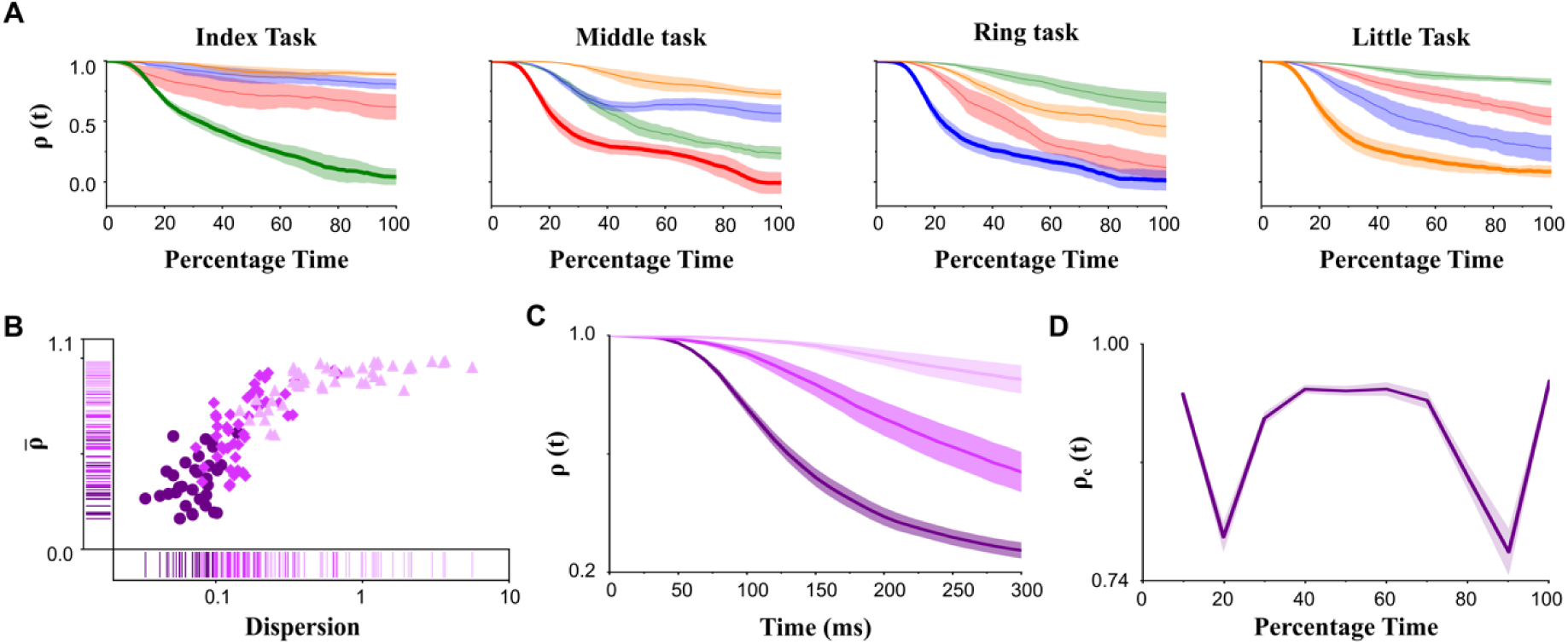
Spearman’s correlation analysis of goal (bold) and non-goal finger movements. A) The trajectories of the correlation coefficient, ρ(*t*), of the four fingers during the goal-directed index finger task, the middle finger task, the ring finger task, and the little finger task (mean ± SEM). The decrease in ρ(*t*) was significantly higher in the goal finger in all four tasks. Colors: green - index, red - middle, blue - ring, and orange - little fingers. B) Plot of average decorrelation (ρ̅) vs trajectory dispersion showed that the goal fingers (circles) had the lowest ρ̅ and lowest dispersion followed by that of non-goal neighboring fingers (diamonds) and non-neighboring (triangles) fingers. C) The trajectories, ρ(*t*) (mean ± SEM), of goal finger, and non-goal neighboring and non-neighboring fingers plotted from movement onset to 300ms into the movement, and the online control was initiated at 60.31 ± 1.93 ms, 115.25 ± 15.13 ms, and 163.64 ± 11.67 ms (mean ± SEM), respectively. D) Correlation analysis of goal finger between consecutive 10 % time instances (ρ_*c*_(*t*), mean ± SEM). The two distinct phases of control, early control at 20 % and late control at 90 % of movement duration, were observed. Colors in Fig. B to D: dark, medium, and light shades of purple represent goal, non-goal neighboring and non-goal non-neighboring fingers, respectively.

**Figure 4:**
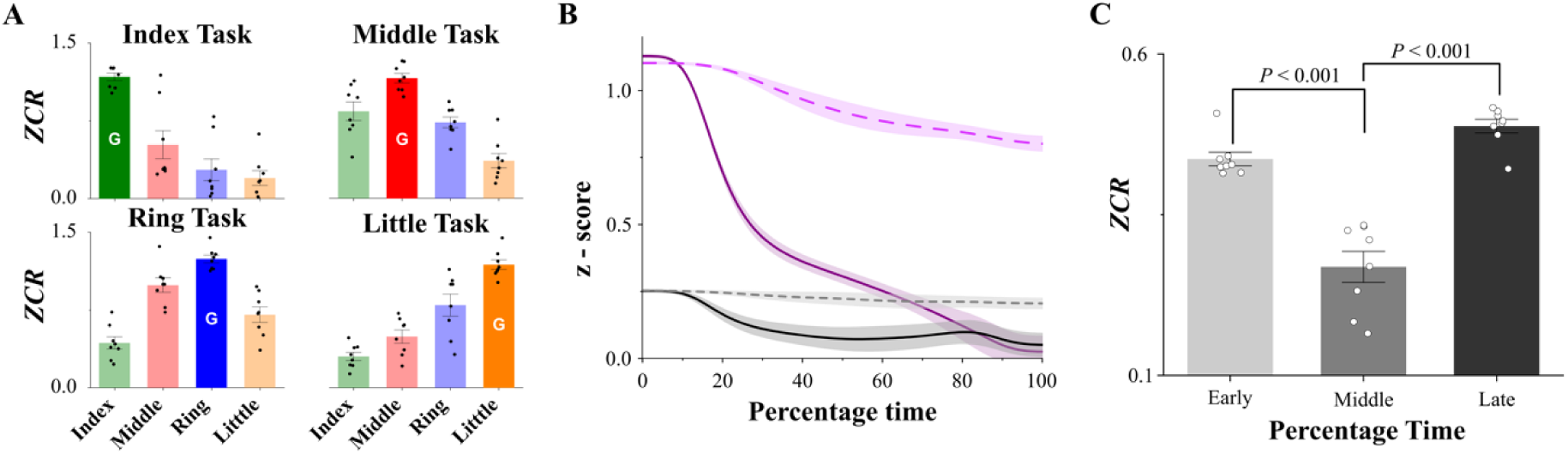
The zero-crossing rate (*ZCR*) of goal (bold) and non-goal finger movements and the average z-scores. A) *ZCR* computed from *z*-scores of goal and non-goal fingers (mean ± SEM) during goal-directed index, middle, ring, and little finger tasks. *ZCR* was the highest in the goal finger of any task (further details in the Results, letter G on the bar represents the corresponding goal finger during any task). Colors: green - index, red - middle, blue - ring, and orange - little fingers. B) *z* - scores of goal (solid line) and non-goal (dashed line) fingers for trajectories (mean ± SEM) starting away (*|z|* > 0.5, dark and light purple respectively) and closer (*|z|* <= 0.5, black and grey respectively) from the average initial posture (*z* = 0). The trajectories that started farther away from the average trajectory had a greater propensity to come towards the average trajectory (*P* < 0.001), which is a potential signature of trajectory control. This convergence towards average trajectory was prominent among goal movements compared to non-goal movements (*P* < 0.001). C) *ZCR* of goal fingers plotted for early (0 to 33 %, light grey), middle (34 to 66 %, medium grey), and late (67 to 100 %, dark grey) movement phases (mean ± SEM). We found that the *ZCR* was significantly greater in the early and late phases as compared to the middle phase (Adj. *P* < 0.001, both cases).

### Experiment 2,3 – Discrete horizontal reaching movements by dominant and non-dominant arms and fast and slow movements

#### Participants

Twenty-five healthy right-handed subjects (15 subjects for experiment 2, and 10 subjects for experiment 3; 21 males; Age: 21-32) participated in the study. The experiment was approved by the institute ethics committee at IISc Bengaluru, and all the participants provided written informed consent before the start of the experiment. The Edinburgh Handedness Inventory (Oldfield, 1971) was used to evaluate their dominant arm.

#### Experimental Setup

Subjects sat on a chair with a backrest to minimize the movements of the clavicle (collarbone). Their dominant or non-dominant arm was required to rest on a smooth elevated table to facilitate horizontal reaching movements (Fig. 1B, left). They were mounted with five electromagnetic 3D motion tracking sensors (Teardrop Mini sensors, 240Hz, Liberty tracking system, Polhemus Inc., Colchester, USA) on the surface of the skin at the sternoclavicular joint (neck), right or left acromion process (right or left shoulder) depending on the task of interest, lateral epicondyle (lateral aspects of the elbow), dorsal tubercle of radius (wrist) and the dorsal aspect of the hand respectively, by a 3M micropore surgical tape. The sensor on the dorsal side of the hand was used for the current analysis. The lights were switched off in the experimental room to occlude the subject’s vision of their arm posture during the movement. Subjects were required to hold an endpoint robotic manipulandum (KINARM, 240 Hz, BKIN Technologies Ltd., Kingston, Canada) and perform a center-out discrete horizontal planar reaching movement as required by the task. During the reach movement, subjects were required to look at the stimulus screen (LCD Screen, 60 Hz) in front of them, where a circular-shaped cursor on the screen was mapped onto the position of the held manipulandum. All the experimental protocols were coded and executed on TEMPO/VideoSYNC Software (Reflective Computing, St. Louis, MO, USA). The dataset was post-processed using MATLAB interface (MathWorks, Massachusetts, USA).

#### Task Structure

A white fixation box (hollow square, 2 cm) appeared at the center of the workspace where the subjects were required to position the pink-colored cursor inside it by moving the manipulandum with one of their arms (Fig. 1B, right). Once they fixated at the center for 500 ms, a red-colored target (filled square, 1 cm) appeared on the screen at an eccentricity of 12 cm from the center at any of the eight directions (0^0^, 45^0^, 90^0^, 135^0^, 180^0^, 225^0^, 270^0^, or 315^0^). Subjects made smooth, unperturbed goal-directed movement towards the target location as soon as the central fixation box disappeared, which was after a hold time of 1000 ms. After initiating the movement, they were given 1000 ms to reach the target location (within a 2 cm square window). Once they reached the target, they were required to stay there for 500 ms to complete one trial successfully. The successful completion of the trial was indicated to the subjects by providing an auditory tone. The inter-trial interval was set to 1500 ms, where the cursor had to be brought back to the central fixation box for the initiation of the subsequent trial. These were called return movements and were not associated with either the primary task or reward. Sample movement traces of goal and return movements by the dominant arm and non-dominant arms are shown in Fig. 1B. If the subject did not fixate enough at the center or at the target, or if they were not able to execute movements within the required time, the trial was aborted. The experiment was conducted such that ∼40 successful trials were recorded in each of the eight directions (5 out of 15 subjects performed ∼60 trials towards 0^0^ and 45^0^, and ∼20 trials towards other directions). The same task was now repeated with the other arm, and the sequence of recruitment of the arms was randomized across subjects.

<u>Experiment 3</u> on fast and slow movements was designed similarly with some minor modifications. The target color was used to cue the subjects about the speed of movement – green for fast (reach within 400 ms) and red (reach from 600-1000 ms) for slow movements, and that the trials were randomly interleaved. For this experiment, the subjects used their dominant right arm and performed movements toward 0^0^ and 45^0^ directions only.

#### Data pre-processing

The motion data from the 3D motion tracking sensors on the hand was low-pass filtered with a cut-off frequency of 20 Hz using a fourth-order zero-lag Butterworth filter (Scholz *et al*., 2000). The time at which the hand crossed 5 % of its maximum speed during the acceleration and deceleration phases, respectively, was noted (Georgopoulos *et al*., 1982; Scott *et al*., 2001; Chakrabhavi & Skm, 2019). The time of onset and the end of movement were identified by descending along the velocity curve from the 5 % mark to whenever it reached zero for the first time. The motion data was then resampled to 100 equally spaced segments to normalize for time (percentage time) across various trials in the analysis of Figs. 5 – 9, while the data for the first 500 ms during the movement was used for the analysis of the actual time scale in Figs. 5C and 7D. For Figs. In 5A, 6A, and 6D, the fixation condition was defined as the time between movement onset (t = 0) to 1000 ms prior to the onset, in the negative direction of time, when the cursor was inside the fixation box. We used this as a reference to compare goal-directed movements that were performed subsequently.

**Figure 5:**
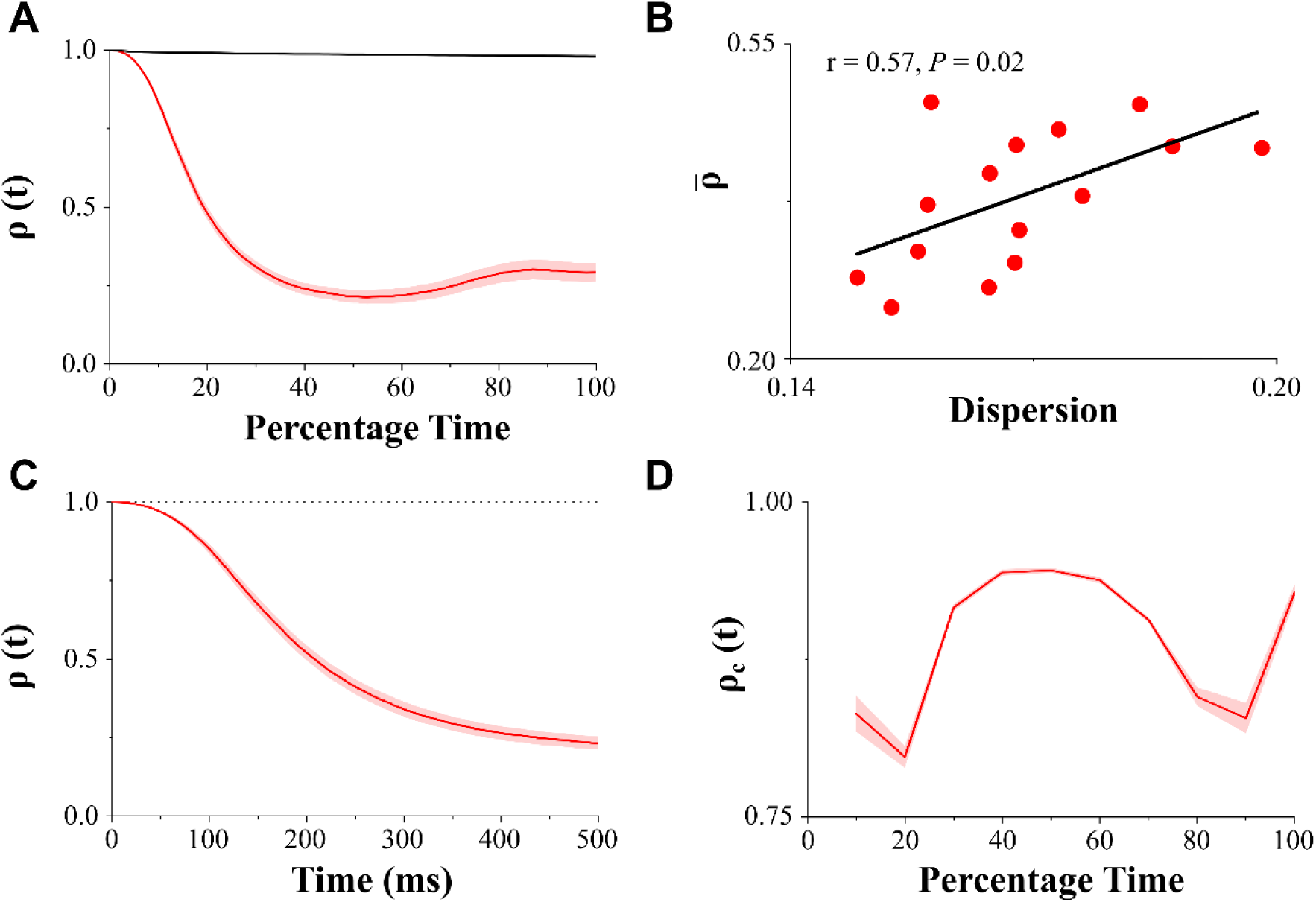
Spearman’s correlation analysis of goal-directed arm movements. A) The correlation co-efficient, ρ(*t*), plotted (mean ± SEM) for goal movements (red) was found to decrease along the movement duration while it stayed close to 1 during fixation condition (black). Goal movements were directed towards the target while the fixation was the time from movement onset to 1000 ms prior to the onset (in the negative direction of time) while the cursor was inside the fixation box. B) The average decorrelation (ρ̅) was significantly associated with trajectory dispersion (Pearson’s r = 0.57, *P* = 0.02). A lower correlation value corresponded with a lower trajectory dispersion. C) The correlation co-efficient, ρ(*t*), plotted on actual time scale (mean ± SEM) for goal (red) movements. The time at which a tangent line (see methods) from ρ(*t*) of goal movements intersected with the ρ = 1, dashed black line was at 69.56 ± 3.45 ms (mean ± SEM). D) Correlation analysis of goal movements between consecutive 10 % - time instances (ρ_*c*_(*t*), mean ± SEM). The two distinct phases of control i.e., early control at 20^th^ percentile and late control at 80^th^ and 90^th^ percentile along the movement duration were observed.

**Figure 6:**
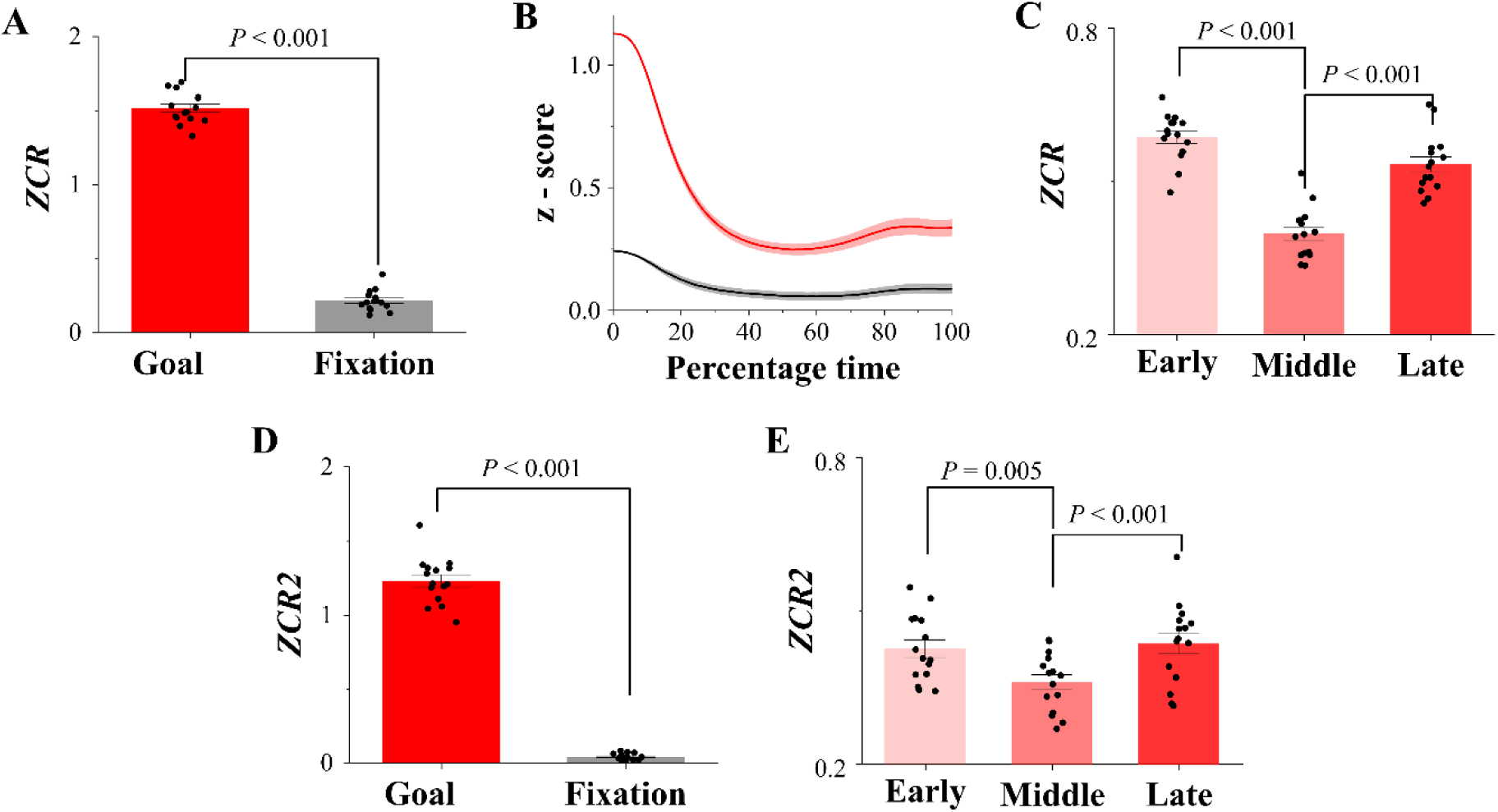
The zero-crossing rate (*ZCR and ZCR2*) and the average z-scores of goal-directed arm movements. A) The 1-dimensional *ZCR* (mean ± SEM) of goal-directed movements (red) was significantly higher than during fixation (black, *P* < 0.001). B) The goal movements were segregated into ones starting farther away (*|z|* > 0.5, red) and closer (*|z|* <= 0.5, black) to the average trajectory (*z* = 0) and we found that the trajectories starting farther away had greater propensity towards the center than the closer ones (mean ± SEM). C) *ZCR* (mean ± SEM) of goal movements computed at the early (0 to 33 %, light red), middle (34 to 66 %, medium red), and late (67 to 100 %, dark red) movement phases. The *ZCR* was significantly greater during early and late phases as compared to the middle phase of movement (Adj. *P* < 0.001, both cases). D) The 2-dimensional *ZCR2* (mean ± SEM) of goal-directed movements (red) was significantly higher than during fixation (black, *P* < 0.001). E) The *ZCR2* (mean ± SEM) was also significantly greater during the early and late phases, compared to the middle phase (Adj. *P* = 0.005 and Adj. *P* < 0.001, respectively).

**Figure 7:**
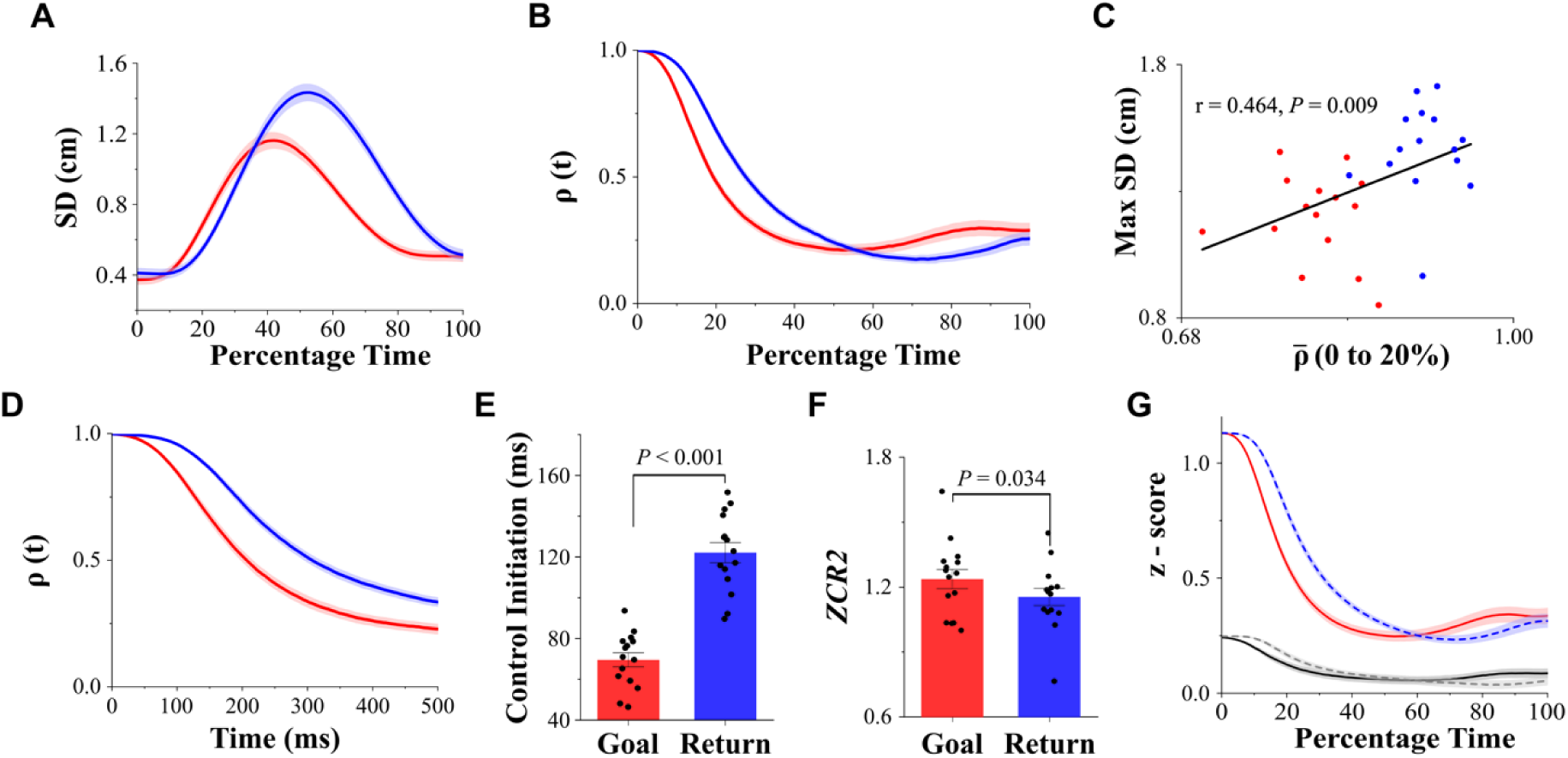
Differences in control between goal (red) vs return (blue) reach movements. A) The trajectory of inter-trial standard deviation (mean ± SEM) for goal and return movements. Return movements had greater SD than the goal movements. B) The trajectories of the correlation coefficient, ρ(*t*), of goal and return movements (mean ± SEM). The average decrease in correlation (ρ̅) was significantly higher in goal movements (*P* = 0.04). C) Correlation between ρ̅, computed from 0 to 20 % time, with the maximum inter-trial standard deviation of goal and return trajectories. The initial decorrelation among trajectories was significantly associated with the maximum inter-trial variability (*r* = 0.464, *P* = 0.009) and that the goal movements had lower ρ̅ and lesser variability than return movements. D) The variations in the correlation coefficient ρ(*t*) on the actual time scale from movement onset to 500ms into the movement (mean ± SEM), showed a steeper and earlier decrease for goal as compared to return movements. E) The initiation of control, marked by the tangent method, showed a significantly earlier onset of control during goal movements as compared to return movements by 52.58 ± 4.75 ms (mean ± SEM, *P* < 0.001). F) *ZCR2* (mean ± SEM) of goal movements was significantly higher than that of return movements (*P* = 0.034). G) The convergence of z-scores (mean ± SEM) towards the average trajectory in the farther away group was significantly greater among goal movements (solid line, red) as compared to return movements (*P* < 0.001, dashed line, blue). The solid black line and dashed grey line belong to the closer group of goal and return movements, respectively.

#### Experiment 4 – Saccadic Eye Movements

The experiment was performed by Varsha et al. (2021) at IISc Bengaluru, and the data have been used for this study as well. A total of 20 healthy subjects participated in this study. Participants sat in front of a monitor, with their heads fixed (Fig. 1C, left), and their eye movements were recorded using an eye tracker (240 Hz, ISCAN, Boston, USA). The experiment was approved by the ethics committee at IISc Bengaluru, and participants provided informed consent.

On a given trial, the subjects had to fixate at the central fixation box for 400 ± 60 ms after which the box disappeared, and a green target appeared in the periphery. The subjects were asked to initiate saccadic eye movements towards the target as soon as possible and received a green tick and a beep sound on successful trial completion. The targets were shown at 12^0^ eccentricities at various orientations (0^0^, 30^0^, 45^0^, 60^0^, 90^0^, 120^0^, 135^0^, 150^0^, and 180^0^) and eccentricities of 6^0^, 8.5^0^, and 10.4^0^ along the orientations of 0^0^ and 90^0^. The saccadic eye movements of 12^0^ eccentricities were considered for the analysis (sample trajectories in Fig. 1C, right). The onset and the end of movement were adopted as it is from Varsha et al., (2021).

#### Data analysis

The displacement data (in the direction of flexion-extension) of the four fingers (index, middle, ring, and little) from Experiment 1 (Chakrabhavi & Skm, 2019), the displacement in X and Y directions (horizontal movement) of the hand from Experiments 2 and 3, and the displacement in X and Y directions of saccadic eye movements in Experiment 4, were used for the analysis.

The number of trials in each of the fingers was selected such that the initial postural variability was equated between the goal and the corresponding non-goal fingers. For the goal and return movements by the arm, the initial variability in hand postures was equalized along both X and Y coordinates. For dominant and non-dominant arm movements, the equalization of initial variability was performed by considering the similarity of movements with respect to both arms. For example, the initial variability along 0^0^ directed movements by the dominant arm was equated with the 180^0^ directed movements performed by the non-dominant arm. The initial variability was similar for fast and slow movements as the trials were randomly interleaved. A standard F-test was used to compare variability between two conditions/distributions, and the farthest point from the mean of the distribution, with more significant variability, was eliminated iteratively until there were no significant differences in variability between the two distributions (Gopal *et al*., 2017). The equalization of initial variability was done to eliminate the influence of initial variance on the correlation analysis and restricted to reporting differences in correlation due to differences in movement execution. For any given task, and at every normalized time point, each trial was assigned a *z*-score using Eq. 1, by subtracting its amplitude from the mean and dividing it by the inter-trial standard deviation (Fig. 2). Subtraction from the mean was performed to track how the trajectories across different trials moved relative to the average trajectory, normalized by the standard deviation, through the course of movement (Figs. 7A and 8A). A sample of movement trajectories (grey) of the arm, directed towards 0^0^, is shown in Fig. 2C, and the *z*-scores of the corresponding red and blue trajectories are shown in Fig. 2D. The motivation of this procedure was to analyze a unitless measure to study variability.

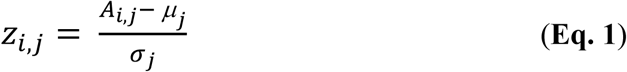

**Figure 8:**
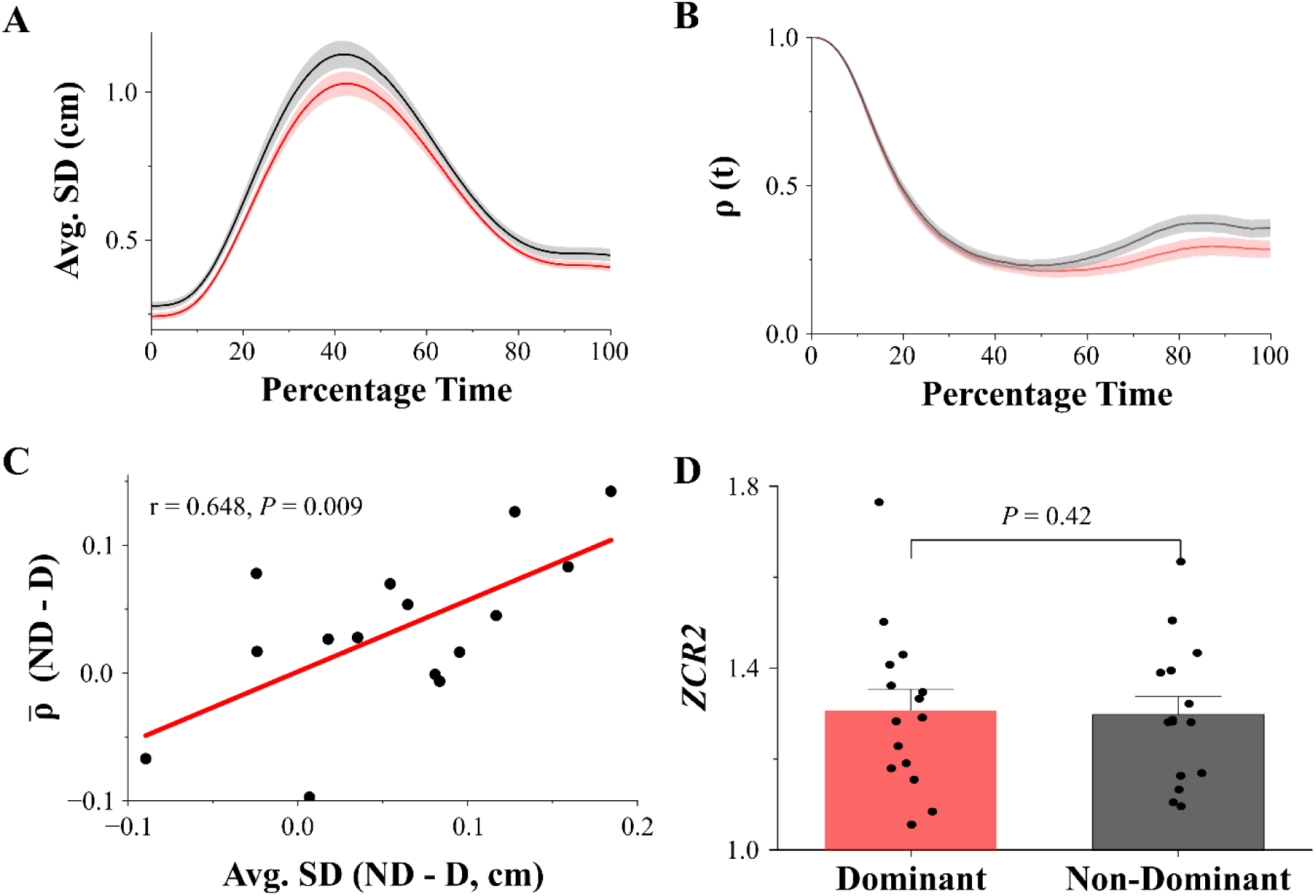
Differences in control between dominant (red) and non-dominant arms (black). A) Plot of inter-trial standard deviation (mean ± SEM) of trajectories of dominant and non-dominant arms. The non-dominant arm had a higher average SD than the dominant arm (*P* = 0.004). B) Correlation coefficient, ρ(*t*) (mean ± SEM), of dominant and non-dominant arms. The average decorrelation was greater in the dominant arm as compared to the non-dominant arm (*P* = 0.029). C) The differences in inter-trial SDs were significantly correlated with the differences in the average decorrelation between dominant and non-dominant arms (r = 0.648, *P* = 0.009). ND – Non-Dominant, D – Dominant. D) The *ZCR2* (mean ± SEM) measure was not significantly different between the two conditions (*P* = 0.42).

where *z*_*i,j*_ is the *z*-score of trial – *i* at time – *j* of the trajectory with amplitude *A*_*i,j*_. Parameters *μ*_*j*_ and σ_*j*_ are the inter-trial average and inter-trial standard deviation of amplitude (*A*) across *n* trials at time *j*. *A*_*n* × *T*_ is the matrix representing the amplitude of trajectories with *n* trials and *T =* 101, normalized time instances.

To investigate the structure of such variability, we performed Spearman’s rank correlation (Hollander *et al*., 2013) between the *z*-scores at the start of movement *z*(0), with the *z*-scores at subsequent time instances *z*(*j*). In Fig. 2, we have depicted two distinct cases of online control. On the left side (Fig. 2A), the representative trajectories crossover each other such that their relative ranks change during the movement. For example, the rank order of trajectories - red, green, blue, and pink (top to bottom) at time *t* = 0, changes to green, pink, red, and blue at time *t* = *j*. This leads to decorrelation among their ranks and a decrease in Spearman’s correlation coefficient (ρ) because of the implementation of active online control.

Conversely, on the right side (Fig. 2B), the order of trajectories remains the same throughout the movement, and the trajectories never converge towards each other, thereby resulting in high ρ (close to 1) and weaker/lack of implementation of online control. We performed the correlation analysis from movement onset to the end of movement (*t =* 0 to 100 %), to obtain a percentage time varying trajectory of correlation coefficient, ρ(*t*). We chose Spearman’s correlation coefficient as we were particularly interested in the crossovers among trajectories, independent of changes in inter-trial standard deviation (Figs. 7A and 8A) during the movement (Janse *et al*., 2021). This method is independent of whether we compute the z - score of the trajectories or do it on the actual amplitudes. We investigated the average decorrelation (ρ̅) over the entire movement duration to quantify the extent of online control. We then fit a logistic function ρ*fit*(*t*), over ρ(*t*) for each of the subject and used parameter *β* to quantify the rate of decrease of ρ(*t*) using Eq. 2.

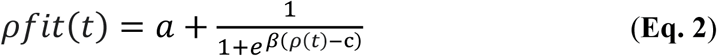

To substantiate our hypothesis, we computed a dispersion measure for the time-normalized trajectories (*A*_*n* x *T*_) by taking the ratio of average inter-trial standard deviation over the time course with the average range of movement during the *n* trials in Eq. 3. We divided variability with the range to account for differences in movement amplitude in the finger experiment. We then correlated the average decorrelation (ρ̅) with its average dispersion measure (or with standard deviation in Figs. 3B, 5B, 7C, 8C, 9C, and 10B), expecting that an increased decorrelation leads to lower dispersion.

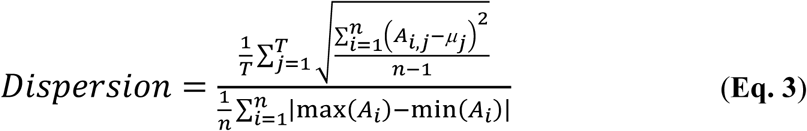

where *μ*_*j*_ is the average of amplitude (*A*) across *n* trials at time *j* and *A*_*i*_ is the trajectory on trial – *i*.

The timing of control in Figs. 3C, 5C, and 7D – E, were done similarly to finding the end of the lag phase and beginning of the growth phase in a growth curve using the ‘tangent method’ (Bertrand, 2019; Smug *et al*., 2024). A tangent was constructed at the point of maximum rate of decorrelation in the Spearman’s correlation coefficient curve, ρ(*t*), and the initiation of control was identified as the time at which the tangent met the ρ = 1, or the horizontal line.

For Figs. 3D, 5D, and 10C, the correlation analysis was performed using a similar technique described earlier but between consecutive 10 % - -time instances. At any percentage time point, the value of Spearman’s correlation (ρ_*c*_) was computed between the *z*-scores of trajectories at that time point with respect to an earlier 10 % - time instance. For example, at *t* = 20 %, correlation analysis was performed between postures at 20 % and 10 % movement duration. This analysis was restricted to goal-directed movements.

We further investigated whether the implemented online control was intended to follow an average planned trajectory. In Fig. 2D, the *z =* 0 represents an average trajectory (dashed line), and the red trajectory crosses the dashed line three times, showing a strong trajectory control. In contrast, the blue line does not cross the dashed line at all, showing weaker trajectory control. To quantify this effect, we computed a zero-crossing rate (*ZCR*) from the *z*-scores of movement trajectories, which was defined as per Eq. 4. A larger *ZCR* implies a greater trajectory control. For arm movements, the *ZCR,* in Figs. 6A and 6C were averaged across X and Y coordinates and along all the movement directions.

The above measure of *ZCR* computed on a one-dimensional dataset was extended to two-dimensional data (ZCR2) in the context of arm movements (Figs. 6D-E, 7F, 8D, and 9D) and saccadic eye movements (Fig. 10D), where trajectories were produced in a spatial plane. This was done by forming a line segment between the trial-averaged 2D spatial locations at any two timestamps t1 and t2 (the average line, the 2D equivalent of a 1D - *z* = 0 line). For any given trial, we approximated the trajectory between the same two timestamps as another line segment. A zero-crossing was marked if the corresponding trajectory line intersected with the average line. Once the number of zero-crossings had been counted for the entire duration and all the trials, *ZCR2* was computed using Eq. 4. We also computed *ZCR and ZCR2* during the early (0 to 33 %), middle (34 to 66 %), and late (67 to 100 %) phases of movements in Figs. 4C, 6C, 6E and 10D.

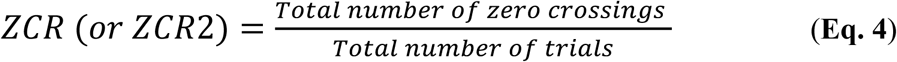

To support evidence of trajectory control, the starting postures of goal and non-goal movements in fingers (Fig. 4B) and arm (Figs. 6B and 7G) were divided into two groups, the ones which were farther away (*|z| >* 0.5, dark and light purple in Fig. 4B, and red and blue in Fig. 6B) and the ones closer (*|z| <=* 0.5, black and grey) to the average initial posture (*z =* 0). The average *z*-scores (across trials) were tracked along the time course of movement to check the propensity of these two groups to converge towards an average planned trajectory (*z =* 0, between 0 – 100 % movement phases). If the central nervous system intended to follow an average trajectory, the group that was farther away from the initial posture should show greater convergence towards the average trajectory than the closer ones. The difference in *z*-scores at 0^th^ (initial) and 20^th^ percentile time points was used to quantify this effect.

### Statistics

The data were checked for normality using the Kolmogorov-Smirnov test. We performed a pairwise t-test for paired comparisons and a two-sample *t*-test for independent comparisons if the data were normal. The Wilcoxon signed rank test and Wilcoxon rank-sum test for paired and independent comparisons were used otherwise. For more than two sample paired comparisons, we performed repeated measures ANOVA. Multiple pairwise comparisons were reported with Bonferroni corrections.

## Results

### Online control of goal and non-goal finger movements

The differences in the magnitude of online control among goal and non-goal fingers were quantified by computing the average decorrelation (ρ̅) during the movement (Fig. 3A). A two-way repeated measures ANOVA (4 finger-tasks x 2 finger-type, goal or non-goal) showed that the ρ̅ of the goal finger was significantly different from the non-goal finger (*P* < 0.001). By comparing the goal and the corresponding non-goal fingers in a pairwise method, it was observed that the extent of decorrelation was highest in the goal fingers (Table 1).

**Table 1:** Pairwise comparison of average decorrelation (ρ^-^) for goal and non-goal fingers (mean ± SEM). The *P* value in the last column is adjusted for multiple comparisons using Bonferroni correction. The ρ̅ The index finger was found to be significantly less than the non-goal fingers.

| Goal Finger | $\bar{\rho}$ | Non-Goal Finger | $\bar{\rho}$ | Adj. *P* |
| --- | --- | --- | --- | --- |
| Index | 0.40 ± 0.05 | Middle | 0.76 ± 0.09 | 0.017 |
|  |  | Ring | 0.89 ± 0.04 | <0.001 |
|  |  | Little | 0.93 ± 0.02 | <0.001 |
| Middle | 0.36 ± 0.06 | Index | 0.56 ± 0.03 | 0.057 |
|  |  | Ring | 0.72 ± 0.03 | 0.003 |
|  |  | Little | 0.87 ± 0.03 | <0.001 |
| Ring | 0.33 ± 0.05 | Index | 0.84 ± 0.04 | <0.001 |
|  |  | Middle | 0.51 ± 0.05 | 0.060 |
|  |  | Little | 0.71 ± 0.04 | <0.001 |
| Little | 0.35 ± 0.04 | Index | 0.92 ± 0.02 | <0.001 |
|  |  | Middle | 0.78 ± 0.04 | <0.001 |
|  |  | Ring | 0.59 ± 0.07 | 0.016 |

The dynamics of online control in goal and non-goal fingers were teased apart by computing the rate of decrease (*β*) in correlation co-efficient, ρ(*t*) (Fig. 3A). A logistic function was fit to the dataset to find *β* (median explained variance = 0.95). A two-way repeated measures ANOVA (4 finger-tasks x 2 finger-types, goal or non-goal) showed that the rate of decrease of ρ(*t*) was significantly different across goal and non-goal fingers (*P* < 0.001). Pairwise comparisons showed that the rate of decrease of ρ(*t*) of the goal-directed fingers was significantly higher than the non-goal, non-neighboring fingers but was not different from that of neighboring fingers (Table 2).

**Table 2:** Pairwise comparison of rate of decrease in correlation (β) for goal and non-goal fingers (mean ± SEM). The *P* value in the last column is adjusted for multiple comparisons using Bonferroni correction. The average rate of decrease in *ρ(t)* in goal finger was found to be significantly higher than the non-goal non-neighboring fingers but was not different from the neighboring fingers.

| Goal Finger | Avg. Rate ($\beta$) | Non-Goal Finger | Avg. Rate ($\beta$) | Adj. P |
| --- | --- | --- | --- | --- |
| Index | $0.077 \pm 0.012$ | Middle | $0.043 \pm 0.024$ | 0.330 |
| | | Ring | $0.030 \pm 0.009$ | 0.028 |
| | | Little | $0.027 \pm 0.006$ | 0.012 |
| Middle | $0.115 \pm 0.032$ | Index | $0.060 \pm 0.013$ | 0.276 |
| | | Ring | $0.057 \pm 0.010$ | 0.255 |
| | | Little | $0.034 \pm 0.006$ | 0.060 |
| Ring | $0.118 \pm 0.032$ | Index | $0.029 \pm 0.004$ | 0.046 |
| | | Middle | $0.078 \pm 0.022$ | 0.230 |
| | | Little | $0.035 \pm 0.005$ | 0.045 |
| Little | $0.131 \pm 0.031$ | Index | $0.024 \pm 0.006$ | 0.005 |
| | | Middle | $0.033 \pm 0.006$ | 0.025 |
| | | Ring | $0.053 \pm 0.016$ | 0.020 |

Further, we checked if the average decorrelation in finger trajectories was systematically associated with the average trajectory dispersion (Fig. 3B) and found that they were tightly correlated (Spearman’s *ρ* = 0.9, *P* < 0.001), and the average dispersion of goal finger (median = 0.073) was significantly less than that of non-goal neighboring (median = 0.143, *P* < 0.001) and non-neighboring (median = 0.573, *P* < 0.001) fingers. Thus, a higher degree of trajectory decorrelation was associated with lesser dispersion in trajectories and vice versa. As goal fingers were associated with varying movement amplitudes, we compared ρ(*t*) of all four goal fingers to check the sensitivity of the correlation analysis to the differences in the range of movement among them. A one-way repeated measures ANOVA on the goal fingers revealed that the ρ̅ were not significantly different (*P* = 0.72), and the rate of decrease of ρ(*t*) were also not significantly different across goal fingers (*P* = 0.52), but the range of movement was significantly different between them (*P* < 0.001), with index (mean ± SEM: 10.63 ± 0.27 cm) and the little (mean ± SEM: 7.42 ± 0.27 cm) finger having the highest and lowest movement ranges, respectively.

Next, we quantified the differences in the initiation of control by performing the correlation analysis on the actual timing using the tangent method. We found that the decrease in ρ(t) of the goal finger (dark purple) was initiated at 60.31 ± 1.93 ms (mean ± SEM). In comparison, decorrelation of the non-goal neighboring (medium purple) and non-neighboring (light purple) fingers were initiated at 115.25 ± 15.13 ms and 163.64 ± 11.67 ms, respectively (Fig. 3C).

Motivated by the previous findings, we examined whether control was constant or occurred during distinct phases along the movement duration. For this, we performed the correlation analysis on the goal finger movements on consecutive 10 % - time instances and found that the controller acted discretely with two prominent troughs in ρ_*c*_(*t*) at 20^th^ (binominal *P* < 0.001) and 90^th^ percentile time (binominal *P* < 0.001) points (Fig. 3D), suggesting the presence of early and late phases of control.

### Trajectory control of goal-directed finger movements

We tested the zero-crossing rate (*ZCR*) of goal and non-goal fingers during the four different instructed finger tasks to probe the propensity of trajectories to follow an average trajectory (Fig. 4A, and Fig. 2D for the computation method). A two-way repeated measures ANOVA (4 finger-tasks x 2 finger-type, goal or non-goal) revealed that the *ZCR* of the goal finger was significantly different from that of non-goal fingers (*P* < 0.001). By performing a pairwise comparison between goal and non-goal fingers, it was found that the goal finger had the highest ZCR in the four tasks (Table 3).

**Table 3:** Pairwise comparison of zero crossing rate (*ZCR*) of goal and non-goal fingers (mean ± SEM). The *P* value in the last column is adjusted for multiple comparisons using Bonferroni correction. The *ZCR* in goal finger was found to be significantly higher than the non-goal fingers.

| Goal Finger | Zero Crossing Rate | Non-Goal Finger | Zero Crossing Rate | Adj. P |
| --- | --- | --- | --- | --- |
| Index | 1.18 ± 0.04 | Middle | 0.52 ± 0.13 | 0.004 |
|  |  | Ring | 0.28 ± 0.11 | <0.001 |
|  |  | Little | 0.20 ± 0.07 | <0.001 |
| Middle | 1.16 ± 0.05 | Index | 0.84 ± 0.09 | 0.045 |
|  |  | Ring | 0.74 ± 0.05 | 0.002 |
|  |  | Little | 0.37 ± 0.07 | <0.001 |
| Ring | 1.24 ± 0.04 | Index | 0.43 ± 0.06 | <0.001 |
|  |  | Middle | 0.99 ± 0.07 | 0.033 |
|  |  | Little | 0.71 ± 0.08 | <0.001 |
| Little | 1.19 ± 0.05 | Index | 0.30 ± 0.04 | <0.001 |
|  |  | Middle | 0.50 ± 0.06 | <0.001 |
|  |  | Ring | 0.80 ± 0.11 | 0.010 |

To support this result further, we divided the initial postures of goal (solid line) and non-goal (dashed line) fingers into two groups (Fig. 4B), the ones farther away (*|z| >* 0.5, purple) and the ones closer (*|z| <=* 0.5, black) to the average initial posture (*z =* 0) and found that the away group had a significantly greater propensity to converge towards an average trajectory (mean ± SEM: 0.43 ± 0.02) than the closer ones (0.09 ± 0.02, *P* < 0.001). Also, the farther away group had a significantly greater convergence in the goal finger as compared to that in non-goal fingers (0.02 ± 0.01, *P* < 0.001). To substantiate the evidence of a temporal structure of control in Fig. 3D, we computed the *ZCR* of goal finger movements across three movement segments (Fig. 4C): early (0 to 33 %, light grey), middle (34 to 66 %, medium grey), and late (67 to 100 %, dark grey) phases of movement duration and found that the *ZCR* was significantly higher during early (mean ± SEM: 0.44 ± 0.01) and late (0.49 ± 0.01) phases as compared to the middle phase (0.27 ± 0.02; Adj. *P* < 0.001 for both the comparisons).

### Online control of goal-directed whole-arm movements

We investigated whether the correlation analysis performed on finger movements could be extended to whole arm reaching movements and if there were systematic control signatures in goal movements. We performed the correlation analysis (Fig. 5A) and found that the ρ̅ in goal movements (mean ± SEM: 0.38 ± 0.02) was significantly less than that during fixation (0.98 ± 0.003, *P* < 0.001). A logistic function was fit over ρ(*t*) (median explained variance = 0.94) to find the rate of decrease (*β*) in correlation, and it was found to be significantly higher in goal movements (mean ± SEM: 0.143 ± 0.005) relative to fixation (0.063 ± 0.028, *P* < 0.005). Further, the average decrease in correlation in goal movements was significantly associated with trajectory dispersion (Pearson’s *r* = 0.57, *P* = 0.02, Fig. 5B). We estimated the timing at which trajectory correction was initiated in goal-directed movements using the tangent method. We found that it occurred at 69.56 ± 3.45 ms (mean ± SEM, Fig. 5C).

Further, to investigate whether such online control was constant or varied across different phases of movement, we performed the correlation analysis of goal-directed movements on consecutive 10% - time instances, ρ_*c*_(*t*). As was observed in finger movements, we found two distinct troughs at 20^th^ (binomial *P* < 0.001) and around 90^th^ (trough shared between 80^th^ and 90^th^ percentiles, binomial *P* < 0.001 for both time stamps, shared across the participants) percentile time instances, indicating early and late phase of control, respectively (Fig. 5D).

### Trajectory control of goal-directed whole-arm movements

As we did with finger movements, we computed the zero-crossing rate (*ZCR*) for goal movements and compared them during fixation and found that the *ZCR* of goal movements (mean ± SEM: 1.52 ± 0.03) was significantly greater than that during fixation (0.22 ± 0.02; *P* < 0.001, Fig. 6A). Similar to the analysis in the previous section, we divided the initial postures of goal movements into two groups (Fig. 6B), the ones farther away (*|z| >* 0.5, red) and the ones closer (*|z| <=* 0.5, black) to the average initial posture (*z* = 0) and found that the away group had a significantly greater propensity to converge towards an average trajectory (mean ± SEM: 0.57 ± 0.02) than the closer ones (0.12 ± 0.02, *P* < 0.001).

To substantiate the evidence for distinct phases of control, the *ZCR* of goal-directed arm movements was computed across three movement segments (Fig. 6C): early (0 to 33 %, light red), middle (34 to 66 %, medium red), and late (67 to 100 %, dark red) phases of movement duration. We found that the *ZCR* was significantly higher during the early (mean ± SEM: 0.59 ± 0.01) and late (0.53 ± 0.02) phases as compared to the middle phase (0.40 ± 0.01; Adj. *P* < 0.001 for both comparisons), like that of goal-directed finger movements.

The above zero-crossing analysis was extended to two-dimensional data with trajectories in the XY plane by computing ZCR2 (see methods). The results were qualitatively similar to the one-dimensional analysis. We found that the *ZCR2* of goal movements (red, mean ± SEM: 1.23 ± 0.04) was significantly greater than during fixation (grey, Fig. 6D, 0.05 ± 0.01, *P* < 0.001). We further divided the *ZCR2* measure of goal movements into three segments (Fig. 6E): early (0 to 33 %, light red), middle (34 to 66 %, medium red), and late (67 to 100 %, dark red) and found that *ZCR2* during middle phase (mean ± SEM: 0.36 ± 0.02) was significantly less than the early (0.43 ± 0.02, Adj. *P* = 0.005) and late phases (0.44 ± 0.02, Adj. *P* < 0.001).

### Differences in control between goal and return movements

As the differences in control measures between fixation and reaches may be related to differences in postural versus movement control, we tested whether these signatures of active control were different across similar movements with varying contexts. For this, we took advantage of return movements that were made back to the fixation spot. The maximum inter-trial standard deviation of return movements (mean ± SEM: 1.46 ± 0.05 cm) was significantly greater than goal movements (mean ± SEM: 1.17 ± 0.05 cm, *P* < 0.001, Fig. 7A). We found that ρ̅ in goal movements (mean ± SEM: 0.38 ± 0.02) was significantly less than that of return movements (0.41 ± 0.01, *P* = 0.04, Fig. 7B). This difference in ρ̅ was prominent in the first 50% of the movement duration (goal: 0.5 ± 0.02, return: 0.61 ± 0.01, *P* < 0.001). A logistic function was fit over ρ(*t*) (median explained variance = 0.97) to find that the rate of decrease (β) was significantly higher in goal movements (mean ± SEM: 0.143 ± 0.05) than in return movements (0.099 ± 0.006, *P* < 0.001). Also, the ρ̅ in goal movements was significantly associated with their respective trajectory dispersion (Pearson’s *r* = 0.53, *P* = 0.002).

Interestingly, the ρ̅ during the first 20 % - time duration was significantly less in goal movements (mean ± SEM: 0.81 ± 0.01) than the return movements (0.91 ± 0.01, *P* < 0.001) and it was significantly associated with the respective maximum standard deviation (Pearson’s *r* = 0.464, *P* = 0.009), which interestingly occurred much later during the movement (Fig. 7C). This association was conserved for the duration as early as 10 % into the movement execution (Pearson’s *r* = 0.426, *P* = 0.018). These differences between goal and return movements were not because of movement kinematics, as the maximum movement speeds were not significantly different across the conditions (*P* = 0.62). Further, the maximum rate of change in ρ(*t*) for goal movements (mean ± SEM: 128.33 ± 5.62 ms) was significantly earlier than that during return movements (201.38 ± 8.02 ms, *P* < 0.001). Using the tangent method, we found that online control during goal movements was initiated significantly faster than the return movements by 52.58 ± 4.75 ms (Fig. 7D-E, mean ± SEM, *P* < 0.001). We also computed *ZCR2* to find the propensity of the trajectories to follow an average trajectory. We found that the *ZCR2* of goal movements (red, mean ± SEM: 1.24 ± 0.04) was significantly greater than the return movements (1.16 ± 0.04, *P* = 0.034, Fig. 7F). In addition, the farther away group in goal movements (mean ± SEM: 0.57 ± 0.02, Fig. 7G) had significantly greater propensity to converge towards the average trajectory than during return movements (0.33 ± 0.02, *P* < 0.001).

### Differences in control between dominant and non-dominant arm movements

As expected, the trajectories of the dominant arm (average SD, mean ± SEM: 0.63 ± 0.02) had significantly less inter-trial variability (Fig. 8A) than the trajectories of the non-dominant arm (0.69 ± 0.02, *P* = 0.004). Although movement kinematics were not significantly different between dominant (maximum speed, mean ± SEM: 23.38 ± 0.98 cm/s) and non-dominant arms (24.38 ± 0.72 cm/s, *P* = 0.33), the ρ̅ was significantly lesser in the trajectories of the dominant arm (mean ± SEM: 0.38 ± 0.02) as compared that of the non-dominant arm (0.41 ± 0.02, *P* = 0.029, Fig. 8B). Differences in the average SD were also significantly associated with the differences in average decorrelation between dominant and non-dominant arms (Pearson’s *r* = 0.648, *P* = 0.009, Fig. 8C). However, no significant differences in the rate of decorrelation (*β*) between dominant (mean ± SEM: 0.145 ± 0.005) and non-dominant arms (0.151 ± 0.006, *P* = 0.19) was observed. Further, the *ZCR2* (Fig. 8D) was also not significantly different between the dominant arm (red, mean ± SEM: 1.31 ± 0.05) and non-dominant arm movements (1.30 ± 0.04, *P* = 0.42). Taken together, we found greater online control in dominant arm movements, but this did not translate to significant differences in trajectory control.

### Invariance of control across different speeds of movement

We subjected the correlation analysis to different movement speeds, between fast and slow conditions (Fig. 9A) to check how sensitive the correlation analysis was to differences in signal-dependent noise that was expected to be greater for higher movement speeds. As instructed, the maximum speed of fast movements (mean ± SEM: 37.03 ± 0.69 cm/s) was significantly higher than the slow movements (mean ± SEM: 16.58 ± 0.39 cm/s, *P* < 0.001). Correspondingly, the max SD was higher during fast movements (mean ± SEM: 0.93 ± 0.02 cm) when compared to the slow condition (0.82 ± 0.03 cm, *P* = 0.002). Nevertheless, we found that ρ(*t*) for fast and slow movements were largely overlapping and that the ρ̅ for fast movements (mean ± SEM: 0.46 ± 0.04) was not significantly different from that of slow movements (0.48 ± 0.05, *P* = 0.317, Fig. 9B). Further, the average decorrelation was not significantly associated with the max SD across two speeds of movement (Fig. 9C, r = 0.249, *P* = 0.289), Despite that, the rate of decrease in correlation (*β*) being greater for slower movements (mean ± SEM: 0.130 ± 0.008) when compared to fast movements (0.108 ± 0.005, *P* = 0.011). We also computed *ZCR2* measure (Fig. 9D) and found no significant differences across fast (mean ± SEM: 0.92 ± 0.08) and slow (1.06 ± 0.12, *P* = 0.089) movements.

**Figure 9:**
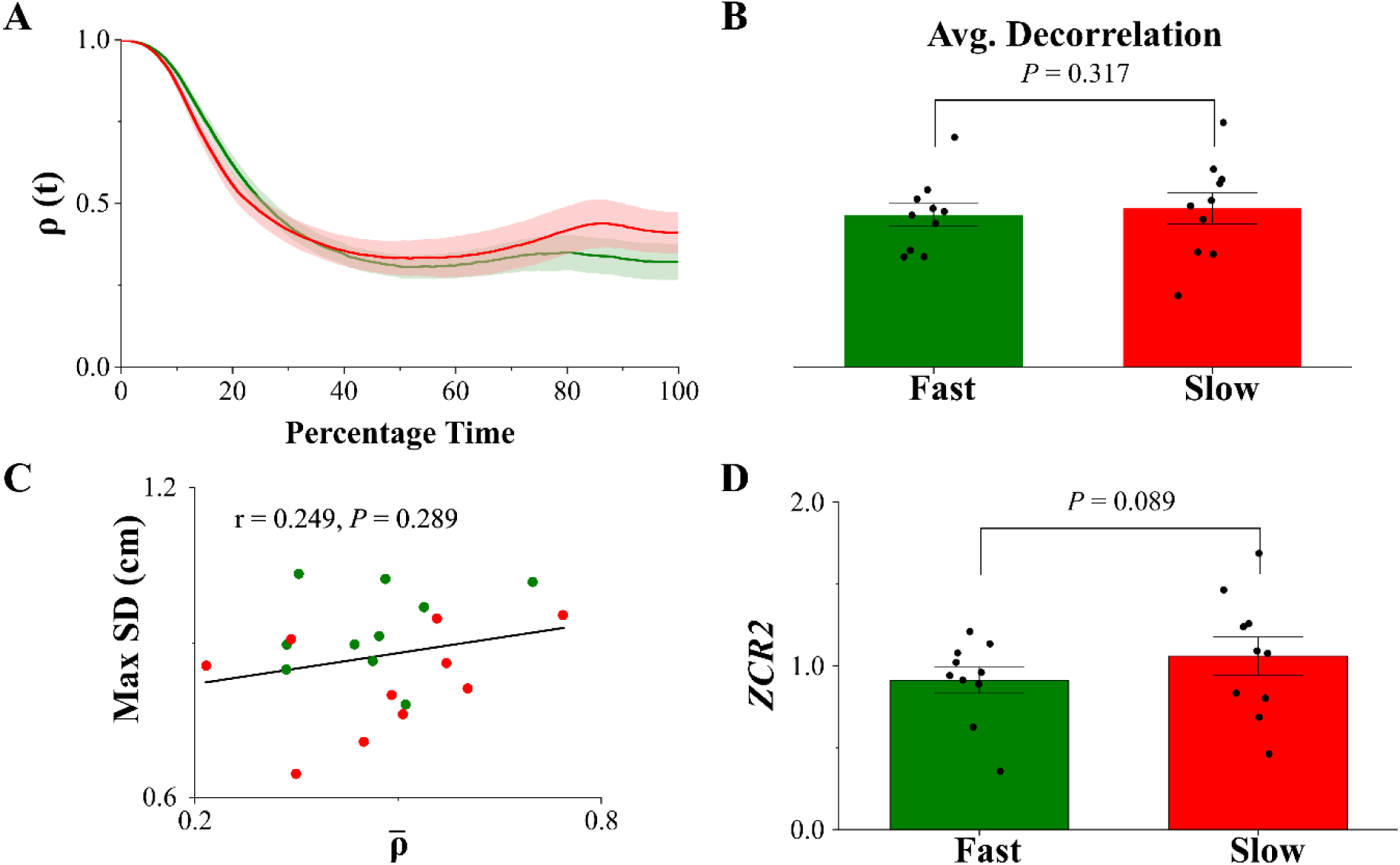
Control analysis on fast and slow reaching movements. A) Plot of correlation coefficient trajectories, ρ(*t*) (mean ±SEM), of fast (green) and slow (red) movements by the dominant arm. The ρ(*t*) of fast and slow movements overlapped with each other. B) The average decorrelation (ρ̅) was not significantly different between fast and slow movements (*P* = 0.317). C) The average decorrelation was not significantly associated with the maximum SD (r = 0.249, *P* = 0.289), even though the maximum SD was higher during fast movements as compared to slow movements (*P* = 0.002). D) The *ZCR2* was not significantly different between fast and slow movements (*P* = 0.089).

**Figure 10:**
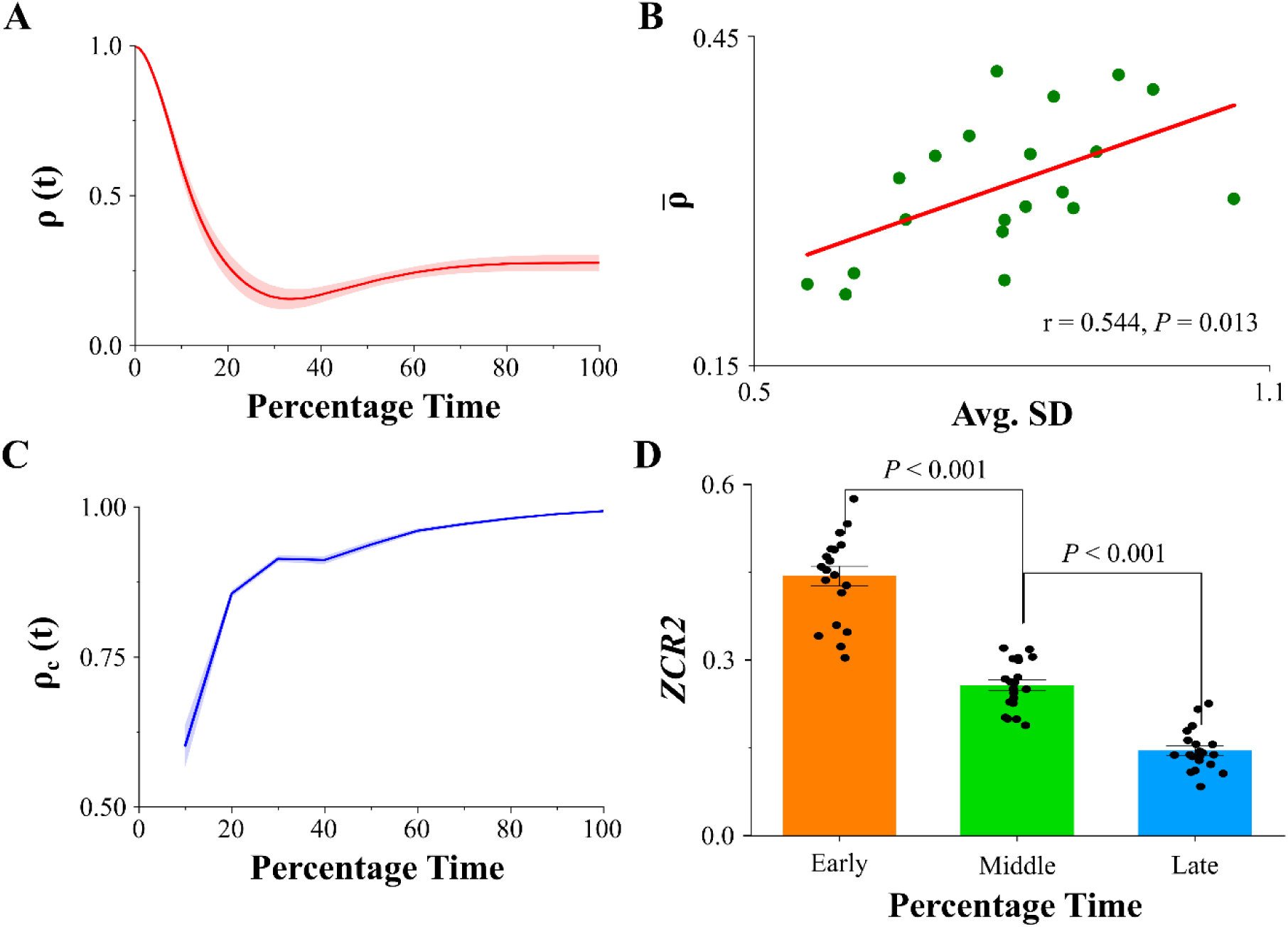
Distinct signatures of control in saccadic eye movements. A) The trajectory of the correlation coefficient (ρ(*t*)) of saccadic eye movements (red, mean ±SEM, averaged across 9 directions). We find signatures of rapid online control with a steep decrease in the correlation coefficient. B) The average decrease in correlation (ρ̅) was significantly associated with the average inter-trial standard deviation (r = 0.544, *P* = 0.013). C) The decorrelation between consequent 10% - time instances (ρ_*c*_(*t*), blue, mean ± SEM) showed it was highest during the early phase of movement and gradually decreased with the time course of movement. This signature of control is unlike that found in finger and arm movements. D) The 2D zero-crossing rate measure (*ZCR2*) of saccades during early (orange), middle (green), and late (sky blue) phases of movement. Trajectory control was the highest during the early phase and decreased along the middle (P <0.001) and late phases (P< 0.001) as seen in Figure C.

### Control of goal directed saccadic eye movements

We subjected discrete goal-directed saccadic eye movements to a similar set of analyses as that of arm movements. We found that the trajectories decorrelated significantly, indicating active online control. The average correlation (ρ̅) was 0.31 ± 0.01 (mean ± SEM), significantly less than 1 (Fig. 10A, *P* < 0,001). The average decorrelation among trajectories was significantly associated with the average inter-trial standard deviation (Fig. 10B, Pearson’s r = 0.544, *P* = 0.013), suggesting that the online trajectory correction was related to a reduction in trajectory variability. The decorrelation analysis on subsequent 10% time instances (ρ_*c*_(*t*)) showed that the saccadic movements did not follow a prominent w-shaped control structure like that of arm movements but rather had control only during the early phase (10% time) of movement (Fig. 10C, binomial *P* < 0.001). Although there was a second significant trough at 40% time (P < 0.001), it was not as prominent as that at 10%. Further, we computed *ZCR2* for early, middle and late phases of movement (Fig. 10D) and found that the trajectory control was predominant in the early phase (mean ± SEM: 0.44 ± 0.02) as compared to middle (0.26 ± 0.01, Adj. *P* < 0.001) and late phases (0.15 ± 0.01, Adj. *P* < 0.001), in support of the previous analysis. It also decreased from middle to late phases of movement (Adj. *P* < 0.001).

Overall, we used the time course of decorrelation between the initial and subsequent time points as a measure of active online control, the rate of zero crossings around a mean trajectory and the trend of z-scores as signatures of trajectory control, and compared these measures in different contexts of finger, arm and saccadic eye movements to show that 1) active online control (or corrections) in trajectories was initiated as rapidly as ∼60 ms in goal-directed finger movements and ∼70 ms in goal-directed whole-arm reach movements, providing behavioral kinematic evidence for fast feedback control during normative movements. 2) Such rapid control also maintained the movements along an average planned trajectory. 3) Control was found to be focused at two distinct phases: at early (20 %) and late (around 90 %) phases along the time course of finger and arm movement. 4) Online control was sensitive to the level of skill (across dominant and non-dominant) but was not sensitive to the range (across instructed fingers) or the speed (across fast and slow) movements. 5) Rapid online control, potentially mediated by internal feedback, was observed in saccadic eye movements, and signatures of control were prominent in the early phase and decreased through the middle and late phases of movement, unlike the w-shaped control structure in the arm movements.

## Discussion

Since Woodworth (1899), the detailed analysis of the kinematics of limb trajectories has provided insights into understanding the relative contributions of motor planning relative to feedback control (Carlton, 1981; Abrams *et al*., 1990; Abrams & Pratt, 1993; Chua & Elliott, 1993). In addition to assessing the ability of the motor system to regulate variability throughout the trajectory (Khan *et al*., 2006), the coefficient of determination (*R*^2^) has been used before as a within-trial estimate of the extent to which early kinematic markers (Carlton, 1981; Gordon & Ghez, 1987; Messier & Kalaska, 1999; Desmurget *et al*., 2005), such as peak acceleration and peak velocity, to determine end-point distributions. In this context, online control has been inferred by measuring the extent to which such markers do not predict the endpoint (i.e., lower *R*^2^). These results, and the validity of the approach, have been validated by manipulating factors such as the availability of visual feedback (Khan & Franks, 2003).

The correlation analysis used in this study uses a similar logic with some important differences that allowed us to reveal novel aspects of control during reaching movements. First, the earliest kinematic markers typically used, peak acceleration at ∼150 ms, are too late to reveal fast feedback control. Here, in contrast to previous works (Messier & Kalaska, 1999; Richardson *et al*., 2011; Kuang & Gail, 2015) that have emphasized the relationship of kinematic markers to the end of movement, we measured how quickly trajectories diverged from an initial starting posture. This analysis ensured that the starting variability between different task conditions was not significantly different and that any difference in control would be revealed at the earliest possible time. Further, the comparison with non-goal movements was particularly useful, since it did not involve manipulation of target locations (Wolpert & Ghahramani, 2000; Gritsenko & Kalaska, 2010) or postures (Pruszynski *et al*., 2008, 2011) that would emphasize differences in sensory feedback control as opposed to internally driven control. Finally, and importantly, the absence of any difference in the extent of decorrelation for fast versus slow movements suggests that signal-dependent noise (Harris & Wolpert, 1998) per se was not a factor contributing to the observed decorrelation signal (Fig. 9).

### Trajectory control during goal-directed arm movements

In contrast to internal models of trajectory control, optimal feedforward/feedback control can also explain straight-line trajectories and is supported by recent work showing the absence of corrective movements to a nominal trajectory in the presence of a perturbation (Cluff & Scott, 2015). Although this result cannot be easily reconciled with our results, we suggest that a potential drawback of perturbation studies to assess trajectory control is the assumption that the original normative plan is still relevant during corrections. In the context of target shift experiments, at least, this assumption is questionable as the original plan is expected to be modified or canceled (Mirabella *et al*., 2008; Murthy *et al*., 2009). By extension, we assume a similar logic would also hold for external perturbations of the hand. Nevertheless, and interestingly, even in the study by Cluff and Scott (2015), when trajectory specifications such as timing and path were part of the goal, corrections were directed to a normative trajectory. In this latter context, our experimental findings broadly agree with them (Cluff & Scott, 2015) as our goal-directed finger and arm movements had timing constraints as compared to the non-goal movements.

Further, the demonstration of relatively fast task-related sensory feedback control (∼50 ms) (Wolpert *et al*., 1998; Pruszynski *et al*., 2008), which could, in principle, shape trajectories during voluntary movement execution involving active participation of the motor cortex (Pruszynski *et al*., 2011), is also a key prediction of optimal feedback control models. However, since movement in optimal feedback models (in the absence of trajectory costs) is driven by costs associated with the current posture and the target position, they don’t necessarily require movements to crossover an average trajectory. This is in contrast with our observations of zero crossings, especially during the early phase, and that the trajectories starting farther away had a rapid propensity to converge towards the average *z* = 0 line (Figs. 4 and 6). This suggests the presence of an average planned trajectory computed from the initial fixation to the target location in the exocentric space. Interestingly, this convergence was prominent for goal movements as compared to non-goal or return movements, implying that they were not a consequence of a regression to the mean effect (Figs. 4B, 7F, and 7G). Furthermore, similar zero crossings across dominant and non-dominant arms (Fig. 8D) and across different movement speeds (Fig. 9D) suggest that the trajectory plan is rather robust across various contexts and is particularly sensitive to goal-directedness.

The other important aspect of control we tested is whether the CNS implements a fixed extent of control or if it varies along different phases of movement. Modeling of perturbation studies on arm movements has suggested that the extent of feedback controller gains is low near movement initiation, while they increase towards the end of the movement as the hand reaches the target (Liu & Todorov, 2007). In the context of finger and arm movements, we found that there were two distinct phases of control (Figs. 3D and 5D), which is reflected at time points where there is a relatively large change of decorrelation in the early (trough at the 20^th^ percentile) and late (trough at 90^th^ percentile) phases. This was also supported by the relatively greater magnitudes of zero crossing rate during early and late phases of movement (Figs. 4C, 6C, and 6E). This pattern is reminiscent of via-point control (Bizzi *et al*., 1984), where, post-perturbation, the arm neither corrected towards the target nor the starting position but ended up at an intermediate point between them. This further reinforces the notion of a trajectory that can be efficiently corrected at two extreme postural positions when the velocities of movements are relatively low and state estimation is more robust. Such distinct troughs were not observed in a previous study since no accuracy constraints were enforced towards the end of movement (Kuang & Gail, 2015). Implementing control close to the target location is obviously beneficial and has been known since the time of Bernstein’s seminal experiments (Bongaardt & Meijer, 2000), and having a high degree of control earlier on has the obvious advantage of clamping variability to manageable levels before the system is swamped by signal-dependent noise. Indeed, we observed that the initial 10 % and 20 % of the response is predictive of the subsequent trajectory variability (Fig. 7C). However, the relative absence of decorrelation between the early and late phases of control suggests that trajectory control is largely intermittent. Thus, we suggest that optimal control models could incorporate such trajectory constraints in their cost function (Mistry *et al*., 2013).

### Trajectory control during goal-directed eye movements

For saccadic eye movements, we found that the trajectory control was predominantly during the early phase (trough at the 10th percentile) and gradually decreased along the time course of movement, suggesting that the structure of control is effectively different from that of arm movements. The association between the average decorrelation and the average inter-trial standard deviation suggests that such decorrelations are likely to be related to the control of these saccadic movements. Although we could not perform the timing analysis due to the lack of enough data points, given that the average duration of saccade was 56.5 ± 2.2 ms (mean ± SEM) and that most of the decorrelation and zero crossings were during the early phase of movement, we suggest that the control could be solely driven by internal feedback consistent with previous work (Guthrie *et al*., 1983; Wurtz, 2008; Richardson *et al*., 2011; Varsha *et al*., 2021).

Interestingly, the absence of the 2^nd^ later peak in saccades relative to arm and finger movements further suggests that such a later phase of control is likely to be mediated by sensory/visual feedback control, which is seen in arm and finger movements and is unavailable in the case of fast saccadic movements. This is also supported by the time evolution of inter-trial variability, which is bell-shaped in arm movements (Figs. 7A and 8A) while it fans out with time in saccadic eye movements (Varsha *et al*., 2021). Nevertheless, in this context, the presence of zero crossings, particularly during the initial phases of the saccadic movement, is consistent with some degree of trajectory control (Vasudevan *et al*., 2023). Although this finding contrasts with standard models of saccadic control that rely on either displacement or end-of-movement (Harris & Wolpert, 1998), it is consistent with recent experimental evidence of trajectory control (Katnani *et al*., 2012; Smalianchuk *et al*., 2018) in monkeys as well as a recently proposed model of saccadic control that simulates kinematic variability in humans during eye-hand coordination tasks (Vasudevan *et al*., 2023).

### Fast feedback control in normative voluntary movements

Several studies have indicated signatures of relatively fast feedback control involving reflexive arm movements that occur between 50 – 100 ms after the beginning of movements during tasks that require posture/ trajectory stabilization following mechanical perturbations (Messier & Kalaska, 1999; Pruszynski *et al*., 2008, 2011; Nashed *et al*., 2014; Cluff & Scott, 2015). However, whether these feedback signals are also at play during unperturbed normative voluntary goal-directed movements is still not clear. In contrast to such fast sensory feedback, internal feedback control has been shown to implement corrections using an efference copy of the initiated motor command to estimate the hand position via a forward model in the absence of delays associated with sensory feedback (Desmurget & Grafton, 2000). To the best of our knowledge, these rapid control strategies have been shown to occur when goals and decisions are changed during movements (Pisella *et al*., 2000; Gaveau *et al*., 2003; Desmurget *et al*., 2005; LoTemplio *et al*., 2023) to help in fast error corrections. In the context of normative movements, such rapid control has been only studied in the context of fast saccadic eye movements using a similar analysis (Richardson *et al*., 2011). Evidence of rapid control has also been shown in isometric force application (Gordon & Ghez, 1987), but it has not been demonstrated in normative voluntary reaching movements.

In our study, the initiation of control was indexed as the time point of significant decorrelation using the tangent method and this occurred as early as ∼60 ms in voluntary finger movements (Fig. 3C), ∼70 ms for voluntary whole-arm reach movements (Figs. 5C and 7D) and predominantly during the early phase of saccadic eye movements. We interpret this rapid control in reaching movements to be a consequence of internal feedback supplemented with fast sensory feedback processes since the onset of goal based decorrelation occurs less than ∼50 ms (movement duration) in the case of saccadic eye movements as well as in some subjects performing the reach movement task (two participants in Fig 7E), which is too fast to allow for fast sensory feedback to operate. Such rapid control in behavior is posited to be mediated even more significantly earlier by the neural substrates, owing to conduction and electromechanical delays (Cavanagh & Komi, 1979; Van Acker *et al*., 2016). However, this rapid onset of decorrelation was typically followed with a later stage that is consistent with fast sensory feedback control with a maximum rate of decorrelation at ∼94 ms for the goal finger and ∼128 ms for goal movements of the arm (Pruszynski *et al*., 2011; Scott *et al*., 2015).

### Distinctions and similarities across finger, arm, and saccadic eye movements

The first experiment (Chakrabhavi & Skm, 2019) required time-synchronized cyclic flexion and extension movements with the goal/instructed finger while it induced involuntary non-goal movements in the neighboring and non-neighboring fingers because of finger enslavement (Häger-Ross & Schieber, 2000; Li *et al*., 2004). In this context, there was a pragmatic expectation of a trajectory controller for the goal-directed finger movements, and we found evidence for the same when compared with non-goal movements in the enslaved fingers. Nevertheless, whether this form of control was specific to cyclic finger movements or whether it generalized to simple reaching movements with the arm (Schaal *et al*., 2004), where it is not clear if the specification of a trajectory is necessary for the control of such movements (Wolpert & Ghahramani, 2000; Todorov & Jordan, 2002; Liu & Todorov, 2007). This was an open question we attempted to address in this study. In this context, a previous study (Sendhilnathan *et al*., 2021) on saccades comparing movements towards a target as a goal, while task-irrelevant movements during the inter-trial interval were considered non-goal, found significant differences in the kinematic variability between goal and non-goal conditions. Neural correlates of such behavioral differences were also observed in the activity of frontal eye field neurons that showed no decorrelation in beta band power prior to non-goal saccades, in contrast to goal-directed saccades. Further, the Fano factor in frontal eye field neurons was much reduced for goal-directed movements compared to non-goal saccades, which correlated with the larger kinematic variability observed for non-goal saccades (Sendhilnathan *et al*., 2021). Such neural and kinematic differences observed between goal and non-goal saccadic movements motivated the current approach to study whole arm reaching movements by considering movements towards a target as goal-directed while return movements during the inter-trial interval, comprising of exertions towards the center, which was not in contention of a reward or completion of a trial, be considered a proxy for non-goal condition. Although the return movements could potentially be guided in anticipation of a reward on the subsequent trial, nevertheless, goal movements in both finger and arm experiments were directed towards completion of the trial and were rewarded, while the non-goal/return movements were not. Moving ahead, it would be interesting to disambiguate the effects of trajectory control from processes mediating the anticipation of reward. Further, the experimental paradigm involving saccadic eye movements (Varsha *et al*., 2021) did not have a hold time. The extent of hold time is seen to have distinct effects on movement planning (Kapoor & Murthy, 2008). We observed a sharp decrease in correlation in this case, unlike a gradual sigmoidal-like decrease in experiments 2 and 3. We speculate that hold time allows precisely timed corrections, unlike the ballistic spontaneous corrections that we see in experiment 4. Despite these differences in the task design, we found distinct signatures of control in finger, arm and saccadic eye movements that could potentially utilize fast internal feedback control to mediate meaningful corrections towards a planned trajectory during the execution of movements.

## Acknowledgements

We would like to thank all the subjects who participated in this study and colleagues in the labs of VSKM and AM for their assistance during the experiments. This research was supported by the Department of Biotechnology – Indian Institute of Science Partnership Program grant (DBT–IISc, *BT/PR27952/INF/22/212/2018*), Intramural funds from IISc (Institute of Eminence grant from the Ministry of Education), and Science and Engineering Research Board grant (SERB, *CRG/2022/000553*). NC was supported by the Prime Minister’s Research Fellowship from the Ministry of Education (PMRF, *PM/MHRD-18-16074.03*).

